# Antimicrobial efficacy of *in-situ* plasma-generated ozone against *Pseudomonas aeruginosa* biofilms in drains and water-submerged surfaces

**DOI:** 10.1101/2023.08.28.553206

**Authors:** Malgorzata Z. Pajak-Zajac, Adam Dowdell, Anthony Buckley, Hugh E. Potts, Andrew Smith, Declan A. Diver

**Author notes:** School of Physics and Astronomy Kelvin Building, University of Glasgow Glasgow G12 8QQ, United Kingdom.

## Abstract

**Aims:** To demonstrate a novel method of gaseous disinfection that can eradicate biofilms inhabiting elusive regions of plumbing systems.

**Methods & Results:** Highly biocidal ozone is generated *in-situ* using the ambient air inside a model sink and drain, via portable plasma discharge, with the plasma device sealing the treated area, ensuring no ozone escape to the external environment. Subsequent bacterial recovery illustrates an approximate bioburden reduction of 5log_10_ for biofilms suspended in the drain, and 6log_10_ for biofilms placed in the sink.

**Conclusions:** Plasma-generated ozone is a safe and effective method for controlling bioburden in periodically wetted, otherwise inaccessible pipework and drains, both above and below the water line.

**Significance and Impact of Study:** The ozone generation system described has potential for controlling healthcare associated infections (HAIs) and infections linked to closed environments, such as cruise ships, from bacteria arising from waste water systems by aerosolization or splash-back. The method has additional benefits provided by an absence of toxic residues after application, reducing risks to users and the waste water system. Cost effectiveness is high, requiring minimal energy input from the electrical supply and only ambient air (no additional feedstock gas is necessary) to generate ozone.

## 1. Introduction

Drains and other (partially or periodically) water-submerged pipework, such as general sanitary plumbing and tap outlets, present a unique challenge to decontamination in that they are both difficult to access and provide ideal conditions for the growth of biofilms (Wingender, 2011; Butler and Upton, 2023). Consequently, reducing bioburden in these contexts can be very difficult, yet it is vital to ensuring a pathogen-free environment for food preparation areas (food factories or commercial kitchens) and immuno-compromised patient care facilities (hospitals and care homes) (Breathnach *et al*., 2012; Kossow *et al*., 2017; Schneider *et al*., 2012; Halstead *et al.,* 2021). The primary risk from contamination via biofilms is the direct expansion of the biofilm itself into areas surrounding the pipework (Bedard *et al*., 2016), with secondary risks arising from aerosolization of the bioburden through splashing or evaporation (Charron *et al*., 2015; Cholley *et al*., 2008; Garvey *et al*., 2016; Gormley *et al*., 2017; Kotay *et al*., 2017; Volling *et* al., 2021). Adverse consequences from transmission of microbes from waste water systems are heightened with the increasing prevalence of multi-drug resistant microorganisms isolated from such systems (Breathnach *et al*., 2012; Hota *et al*., 2009; Weiner *et al*., 2016; Zarb *et al*., 2012; Kizny Gordon *et al*., 2017).

Following outbreaks of *Pseudomonas aeruginosa*, a problematic gram-negative bacteria, in critical care units of hospitals globally, different preventive maintenance protocols have been proposed (Health Technical Memorandum 04-01: Addendum, 2016). These include: UV treatment of the sink drains, re-design of toilet and shower drains, chemical disinfection and thermal disinfection (Gordon *et al*., 2017; Walker and Moore, 2015; Garvey *et al*., 2019). A plasma is a partially ionized gas which has the ability to create ions and free electrons as well as highly reactive oxygen species (ROS) and nitrogen species (RNS) (Lu *et al*., 2016), all of which contribute to the ambient gas chemistry and its associated disinfectant properties. One such ROS that can be generated via plasma is ozone, a well-established biocide that inactivates microorganisms through oxidation of cell wall components (Hirneisen *et al*., 2011; Kaptan *et al*., 2014; Lim *et al*., 2010; Nicholas *et al*., 2013; Saha *et al*., 2014; Zargaran *et al*., 2017, Akter *et al*., 2020). Most of the literature concerning eradication or inactivation of microbial biofilms via plasma, published to date, focuses on the effects on various ROS generated in plasma-activated water, or direct interactions between plasma and biofilms (Huth *et al*., 2009; Nakano *et al*., 2015; Soler-Arango *et al*., 2018; Vandervoort and Brelles-Marino, 2014; Zelaya *et al*., 2010; He *et al* ., 2020). The technique described here is an indirect one: the plasma is generated *in-situ* from the ambient air inside a gas-tight volume via a dielectric barrier discharge (DBD) and is confined to a thin layer at the electrode assembly; the resulting plasma chemistry (including ozone) is distributed throughout the region being treated, such that there is no direct impact of the plasma on the treatment surface. There is no requirement to open the sealed region for access, nor is any feedstock gas required. The aims of this manuscript are to describe the bactericidal effect of ozone generated *in-situ,* via a DBD system, on *P. aeruginosa* biofilms in an in vitro sink drain model.

## 2. Methods

### 2.1 Ozone generation

The patented plasma system (patent EP 2497343 B1 WO 2011/055113) consists of a high-voltage power supply (4-5 kV, 10-50 kHz) fully contained within an electrically grounded shield (Diver and Potts, 2010). Power is supplied to a DBD solid monolithic plasma electrode system in which the powered electrode is entirely internal and separated from the external structured electrode by a layer of insulating material (Fig. 2). Samples to be treated with ozone are placed into generic polyethylene grip-seal bags with the plasma source effectively sealing the mouth of the bag, creating a closed space filled with ambient air; electric field then penetrates the interior of the bag due to the gaps in the outer electrode, ionizing the ambient air inside the bag and producing ozone (Diver and Potts, 2010). It is also possible to produce ozone within a target chamber, such as a drain limited by the water trap on one end and plasma generating device on the other, once again using the ambient air inside (Fig. 3). Given that the ozone yield produced by our source reduces with the rising electrode temperature of the plasma head (data not shown), an infrared thermometer (62 Max, Fluke, Everett, WA, US) was used to monitor the plasma head temperature before and after treatment; this ensured it did not exceed 33°C, such that all samples received comparable doses of ozone.

#### 2.1.1 Ozone environmental monitoring

The experiments were conducted on an open bench. Given the workplace exposure limit (WEL) of ozone (0.2 ppm in air averaged over a 15 min reference period (Health and Safety Executive, 2014)), an ozone personal monitor (BW GasAlert Extreme, BW Technologies by Honeywell, Hegnau, Switzerland) was always placed adjacent to the experiment to ensure no ambient ozone leakage. In addition, for the model wash basin experiment, the output pipe was sealed by a collecting bag, in which a pre-wetted ozone corn starch-iodine indicator strip was placed (MN Ozone, Camlab, Cambridge, UK). No ozone leakage was detected.

#### 2.1.2 Measurement of ozone generated

A generic clear polyethylene grip-seal bag filled with ambient air, in the absence of any substrate, was treated with ozone using the same parameters as the samples. Once the treatment concluded an ozone meter (106-MH, 2B Technologies Inc., Boulder, Colorado, USA) was used to sample the air inside the bag at 10s interval, to measure ozone concentration. The meter was connected to a computer via USB interface and the measurements logged into a text file using custom-written Python software. This allowed for the calculation of maximum ozone concentration and total yield.

### 2.2 Microbial strain and culture media

A *P. aeruginosa* NCTC 10662 was used in all experiments. If not otherwise specified, Tryptic Soy Broth (TSB) (Sigma-Aldrich, Gillingham, UK) was used as a rich nutrient medium, and recovered bacteria were cultured onto Tryptone Soy Agar plates (TSA) (E&O Labs, Bonnybridge, UK).

### 2.3 Biofilm formation on coupons

A CDC bioreactor (BioSurface Technologies Corporation, Bozeman, MO, USA) was used to generate a reproducible *P. aeruginosa* biofilm in high shear conditions. The method is described in ASTM E2562-12 standard (ASTM International, 2012), with the temperature modified in some of the experiments (20-22°C instead of 30°C). The arms of the CDC bioreactor were fitted with ½ inch polycarbonate or polytetrafluoroethylene (PTFE) coupons, and the bioreactor filled with 35 ml of TSB and 315ml sterile deionised H_2_O. It was then inoculated with 1-1.6 ml of *P. aeruginosa* culture (ca. 10^9^ CFU ml^−1^). The biofilm was then grown with agitation (130 rpm) for up to 24h, at 20-22°C. For more mature biofilms, the bioreactor was run for an additional 48h in the continuous phase, which entailed feeding the bioreactor with diluted medium (16.7 ml of TSB per 5,000 ml of sterile deionised H_2_O) at 5 rpm. Each arm (8 total) can accommodate 3 coupons, allowing for a total of 24 biofilms to be cultured. The temperature is kept consistent across all arms for any given experiment. Upon reaching a defined timepoint for growth, arms are removed from the bioreactor and gently submerged in a tube with 50 ml sterile PBS or ¼ strength Ringer’s solution with 0.05% polysorbate 80 to dislodge planktonic cells and provide consistently biofouled coupons.

### 2.4 Method of biofilm recovery by sonication

The ASTM standard E2562-12 was modified by replacing scraping as a method of recovery with sonication. Sonication has been demonstrated to be more reproducible, as well as being the most effective method for removal of biofilms from medical devices (Assere *et al*., 2008; Bjerkan *et al*., 2009; Webber *et al*., 2015, Singh *et al*., 2021). For this method, treated or untreated biofilms, grown on the specified coupon type and under the desired conditions, as required, were aseptically transferred to glass universal containers filled with 10 ml of a sterile neutralizer (PBS or ¼ strength Ringer’s solution with 0.05% polysorbate 80) and sonicated for 10 min in an ultrasonic bath (Medisafe Reliance PC+ 5L, Medisafe UK Ltd, Bishop’s Stortford, UK). The resultant bacterial suspension was then diluted as required, plated on TSA plates, then incubated at 37°C overnight and enumerated. Verification of a *P. aeruginosa* biofilm was confirmed by colony morphology and Gram stain.

### 2.7 Experiments

#### 2.7.1 Ozone vs. control disinfectant

In order to provide a reference benchmark for the antimicrobial activity of the DBD system generated ozone, we compared ozone disinfectant efficacy to a surface liquid disinfectant marketed in Europe for many applications, including healthcare. The liquid disinfectant used was a freshly prepared 1% solution of a Pentapotassium bis (peroxymonosulphate) bis (sulphate) (Rely+On Virkon, DuPont UK Ltd., Bristol, UK).

Biofilm was grown on polycarbonate coupons, as in section 2.3, at room temperature (RT) for 8 and 12h. For both times, bioreactor arms were allocated for coupons to be treated with ozone and disinfectant, as well as a control group – to be untreated. A whole bioreactor arm with three coupons was placed in a generic clear polyethylene grip-seal sampling bag filled with an ambient air; the bag was then subjected to a cold plasma to generate ozone, with plasma generated for 100 s at 4.7 kV, and left for a dwell time of 1h. The untreated samples were handled the same, but without any exposure to ozone.

For samples treated with liquid disinfectant, 10 ml of a freshly prepared 1% solution of Rely+On Virkon (SLD) was aliquoted to three universal containers. A biofilm-covered coupon was added to each container and all containers were subsequently kept in a water bath set to 40°C for 10 min. Afterwards, the coupons were removed and gently rinsed twice in sterile deionised water. The bioburden of each sample was then quantified as in section 2.4. This data is presented in Table 1.

#### 2.7.2 Ozone treatment of biofilms of different maturity

Biofilms were grown as in section 2.3, on PTFE coupons and at RT, with each set of 2 arms in the bioreactor containing coupons to be retrieved after growth timepoints of 12, 24, 48 and 72h. At the specified timepoints coupons were removed aseptically and bagged individually into generic clear polyethylene grip-seal bags. Samples from the first arm in each set were treated using plasma generated *in-situ* for 100s at 4.7 kV. Control samples were left untreated with ozone. Bacterial growth was then quantified as in section 2.4. Results are illustrated in Table 2.

#### 2.7.3 Ozone treatment of a model wash basin and drain

A model wash basin and drain assembly was constructed using commercially available parts, to test the efficacy of plasma generated ozone as a system for decontamination in complex geometries. It consisted of a 290mm stainless steel square mini-wash bowl (no overflow or tap holes and a central drain), a vertical polypropylene outlet pipe ending with a polypropylene bottle trap (Floplast; Ø 40 mm inlet and outlet, compression fittings) and an effluent pipe. The system rested on a speed frame (Rexroth aluminium strut 20mm × 20mm). The bottle trap was filled with fresh tap water just below the overflow level to the effluent pipe. An experiment diagram and a picture of the model wash basin and drain assembly are shown in Fig. 3.

Instead of using a bag as the treatment chamber, the drain was treated with a device that rests on the basin edge, around the plughole. This device consists of the DBD plasma source of the same size and parameters as the one used in the other experiments described in this work, but encased in an oval dome (Potts and Diver, 2015). This design allows for creating a closed region in which ozone could be generated, using the water in the drain trap as an effective seal (Fig. 3). Upon activation of the source, oxygen from the air within the dome is converted into ozone, with the rubber seal around the edge of the dome providing a gas-tight seal. The underlying physics and chemistry are the same as previously described.

The treatment protocol consisted of a series of brief plasma pulses, specifically 30s bursts with subsequent 30s intervals, for a total of 30min, yielding an ozone concentration over time as illustrated in Suppl. Fig. 6. A dwell time of a further 30min ensured that the ozone dissipated completely. The ozone concentration within the dome was measured using an ozone meter as described in section 2.1.2.

A 13h biofilm was grown on several PTFE coupons in the CDC bioreactor, at RT, according to section 2.3. All coupons had a short piece of PTFE tubing (inner Ø 1 mm, 3-4 mm length) attached to one side using silicone and placed in a manner that they faced the bioreactor wall. Three coupons were placed in the model wash basin, spaced equally. Another three were suspended on a nylon filament and hung in the vertical outlet pipe: the bottom one just above the surface of the water, the top one just below the drain and the middle coupon approximately mid-way between the top and bottom coupons, as shown in Fig. 3a. After treatment all coupons were removed, alongside the PTFE tubing and silicone. One set of three sterile PTFE coupons was used as negative control and one set of three untreated coupons from the same biofilm production batch was used as positive control. Bacterial growth was quantified as in section 2.4.

#### 2.7.4 Biofilm regrowth on ozone inactivated biofilm

We investigated whether previously biofouled and ozone treated coupon surfaces have an increased potential for bacterial growth. Biofilm was initially grown on polycarbonate coupons at 30°C for 24h, as described in section 2.3. Of the total 21 coupons (across 7 arms), 18 were put individually in generic clear polyethylene grip-seal bags and treated with ozone as in previous experiments (plasma generated for 100 s, 4.7 kV). Of the treated coupons, 9 had their bioburden quantified according to section 2.4, then were cleaned and autoclaved; the 3 untreated (control) coupons were processed similarly. Those 12 coupons were then fitted back into the bioreactor, along with the other 9 previously ozone treated coupons. The bioreactor was restarted using the same parameters, with the cleaned and ozone treated coupons arranged in opposing arms. Samples were taken from each at 4, 6, 8 and 26h. The produced biofilm was recovered and enumerated as in section 2.4.

## 3. Results

Ozone generated from air via a DBD has an upper concentration limit, where generation and loss are in equilibrium, that is a strongly correlated with humidity; increased humidity has a detrimental effect on ozone production (Ki *et al*., 2019). For the experiments reported here, using ambient air inside a small sealed polyethylene bag, the concentration limit was typically about 1800 ppm of ozone; the time required to generate maximum concentration depends on the bag volume to surface area ratio. Using our technology, around 3 mg of ozone can be produced in a 1-litre bag in 1 min. Typical ozone yields are shown in Suppl. Fig. 1, showing that 30s of operation yields 1.5mg of ozone in the interior of the package, just over 700 ppm for a one litre bag, for example.

The limit of detection (LOD) for biofilm recovery via sonication was calculated based on quantities and dilutions used in a validation of this technique as a method for quantifying bioburden (see Suppl. section 1.3.1). This value was estimated at 50 CFU ml^−1^.

### 3.1 Ozone vs. control disinfectant

The number of recovered total culturable *P. aeruginosa* from the biofilms grown for 8 and 12h on polycarbonate coupons subjected to gaseous ozone and standard, SLD treatment is summarised in Table 1. With both treatments, no culturable organisms were detected, which was interpreted as a log_10_ reduction of at least 3.4 and 4.3 for the 8 and 12hr biofilms, respectively.

### 3.2 Ozone treatment of biofilms of different maturity

The treatment of biofilms grown at RT on PTFE coupons, sampled at 12 to 72h and exposed to ozone, rendered no detectable culturable *P. aeruginosa*, as shown in Table 2.

### 3.3 Ozone treatment of a model wash basin and drain

The bacterial recovery from untreated coupons was (4.4 ± 0.2) × 10^7^ CFU ml^−1^. No culturable organisms were recovered from the coupons in the model wash basin, while the mean number of recovered organisms from the drain was (1.9 ± 1.8) × 10^2^ CFU ml^−1^. This is a log_10_ reduction of at least 5.9 for the wash basin coupons and at least 5.1 for the coupons suspended in the drain. No ozone was detected escaping from the water trap during this experiment.

### 3.4 Biofilm regrowth on ozone inactivated biofilm

The number of culturable organisms recovered from PTFE coupons on which a fresh *P. aeruginosa* growth cycle has been induced, after previous biofilm growth and subsequent ozone treatment, is shown in Figure 1. There was no detectable difference in biofilm regrowth between coupons with previous biofilm growth, and subsequent ozone treatment, compared to previously untreated coupons.

### 3.5 Tables and Figures

**Table 1.**
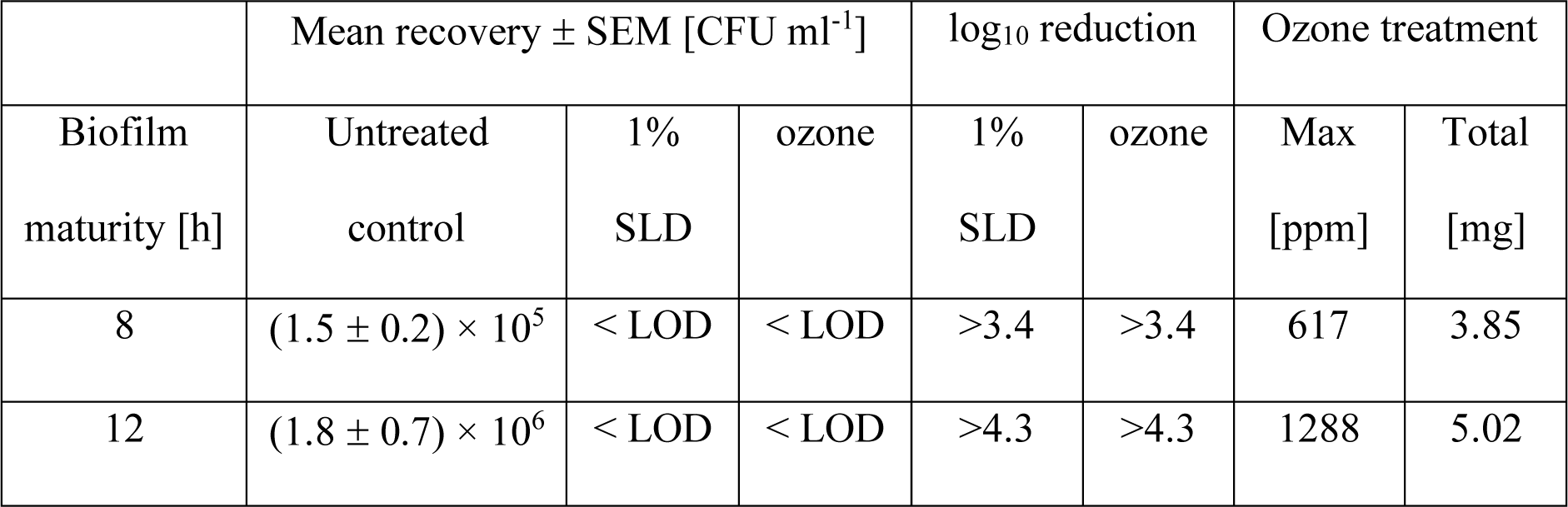
Efficacy of treatment of different maturity biofilm grown on polycarbonate coupons.

| Biofilm maturity [h] | Mean recovery ± SEM [CFU ml<sup>-1</sup>] | | | log<sub>10</sub> reduction | | Ozone treatment | |
| --- | --- | --- | --- | --- | --- | --- | --- |
|  | Untreated control | 1% SLD | ozone | 1% SLD | ozone | Max [ppm] | Total [mg] |
| 8 | $(1.5 \pm 0.2) \times 10^5$ | < LOD | < LOD | >3.4 | >3.4 | 617 | 3.85 |
| 12 | $(1.8 \pm 0.7) \times 10^6$ | < LOD | < LOD | >4.3 | >4.3 | 1288 | 5.02 |

**Table 2.**
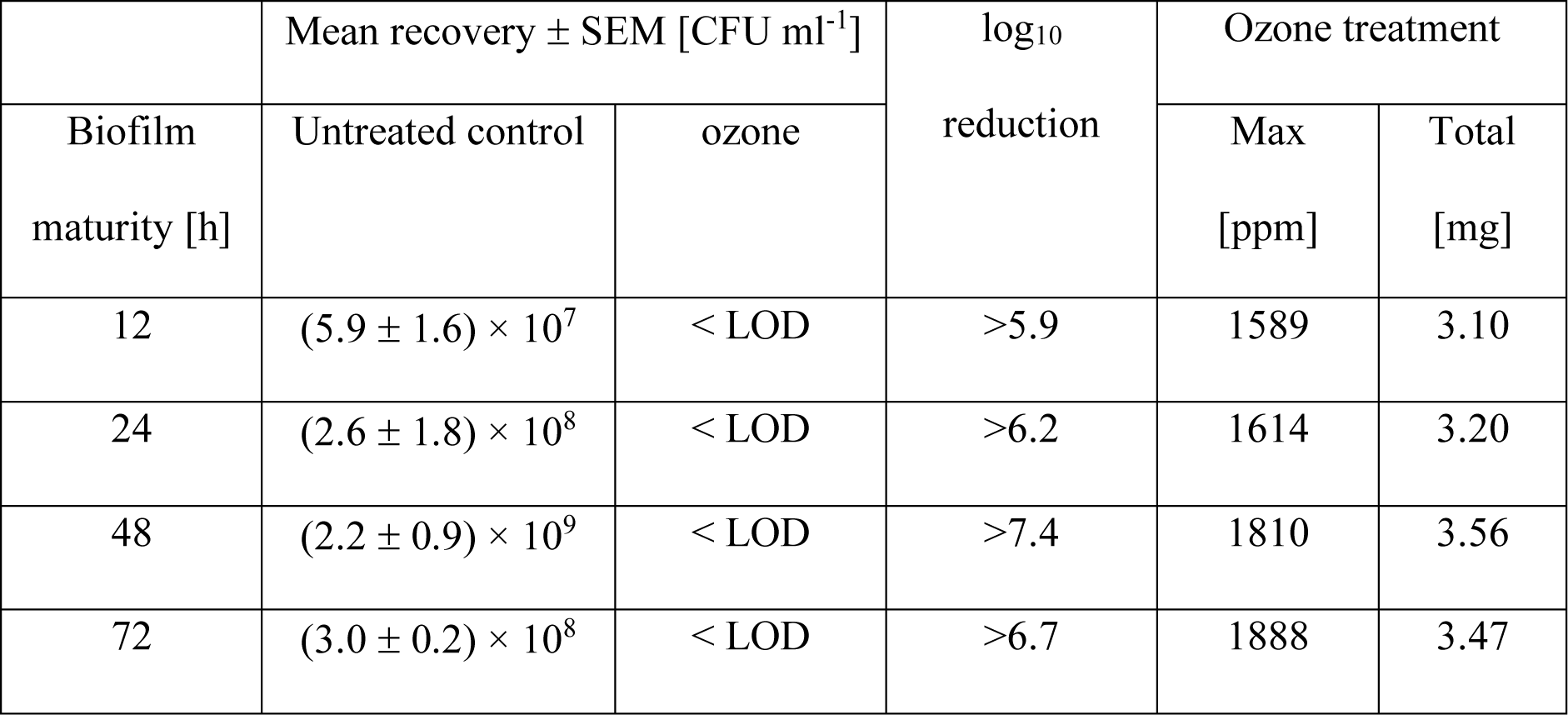
Inactivation of biofilms grown for up to 72h on PTFE coupons.

| Biofilm maturity [h] | Mean recovery ± SEM [CFU ml<sup>-1</sup>] | | log<sub>10</sub> reduction | Ozone treatment | |
| --- | --- | --- | --- | --- | --- |
|  | Untreated control | ozone |  | Max [ppm] | Total [mg] |
| 12 | $(5.9 \pm 1.6) \times 10^7$ | < LOD | >5.9 | 1589 | 3.10 |
| 24 | $(2.6 \pm 1.8) \times 10^8$ | < LOD | >6.2 | 1614 | 3.20 |
| 48 | $(2.2 \pm 0.9) \times 10^9$ | < LOD | >7.4 | 1810 | 3.56 |
| 72 | $(3.0 \pm 0.2) \times 10^8$ | < LOD | >6.7 | 1888 | 3.47 |

**Fig. 1.**
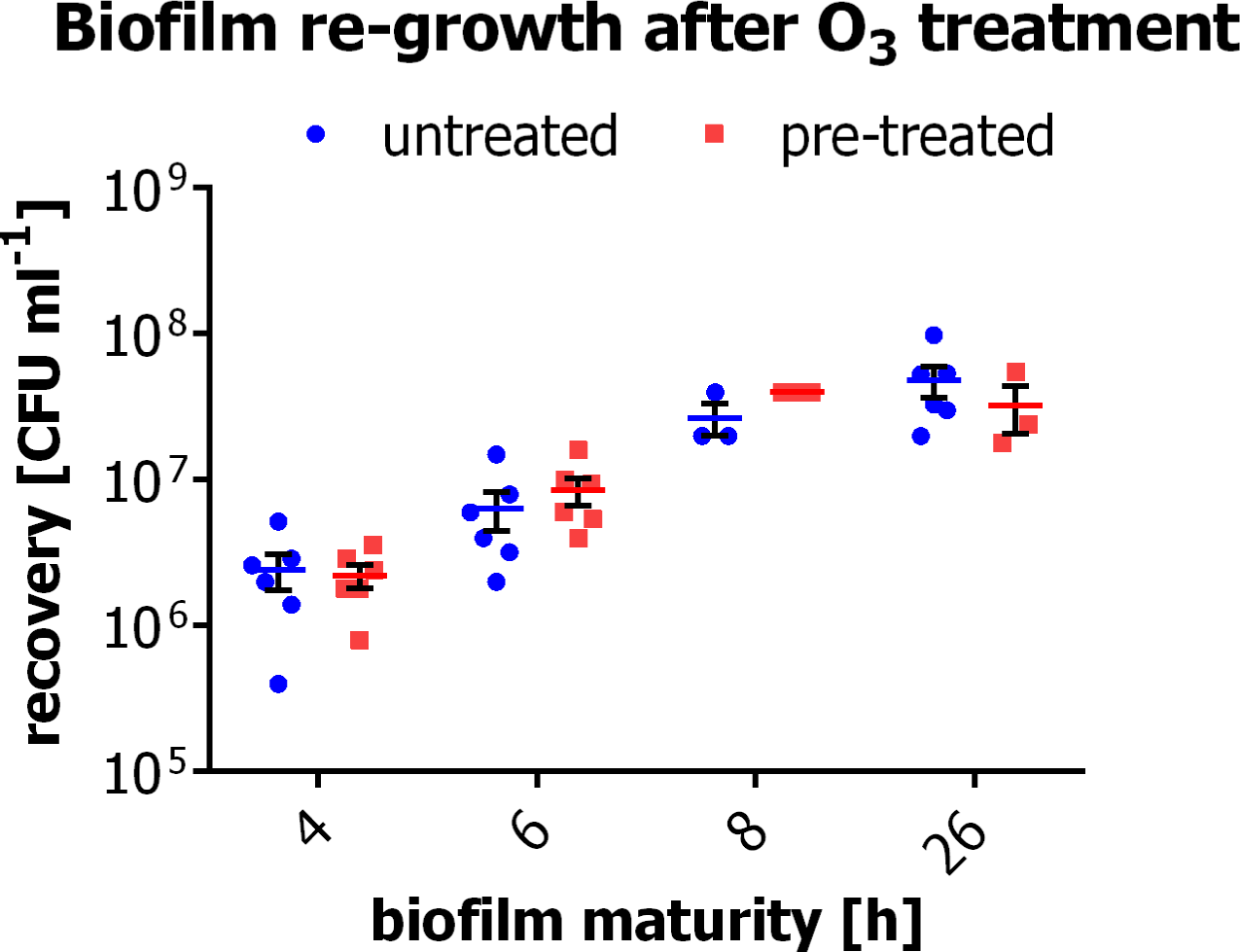
Recovery of total culturable P. aeruginosa after previous ozone treatment. Error bars represent SEM.

**Fig. 2.**
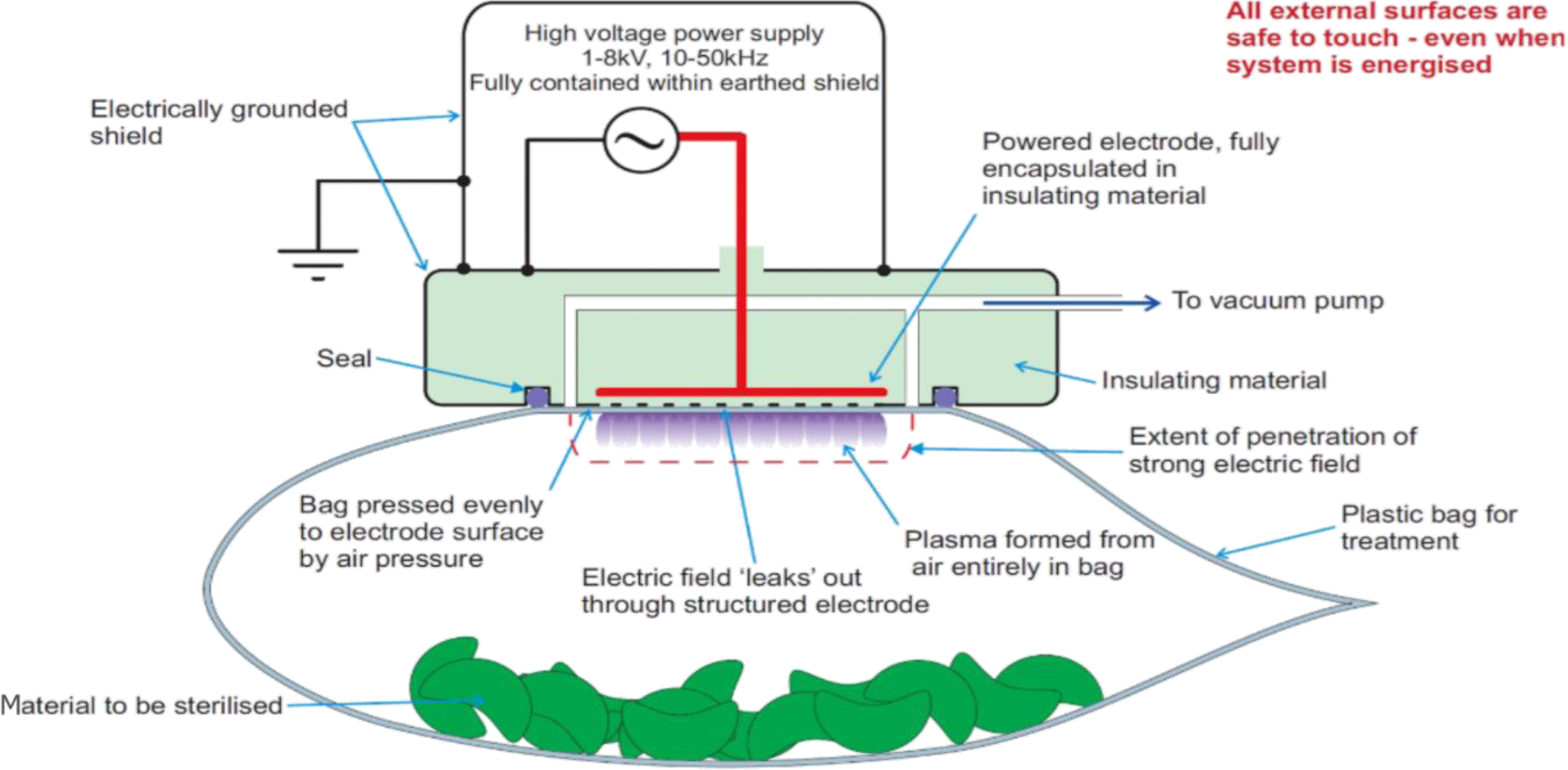
Schematic of plasma DBD system showing electrode arrangement and operation.

**Fig. 3.**
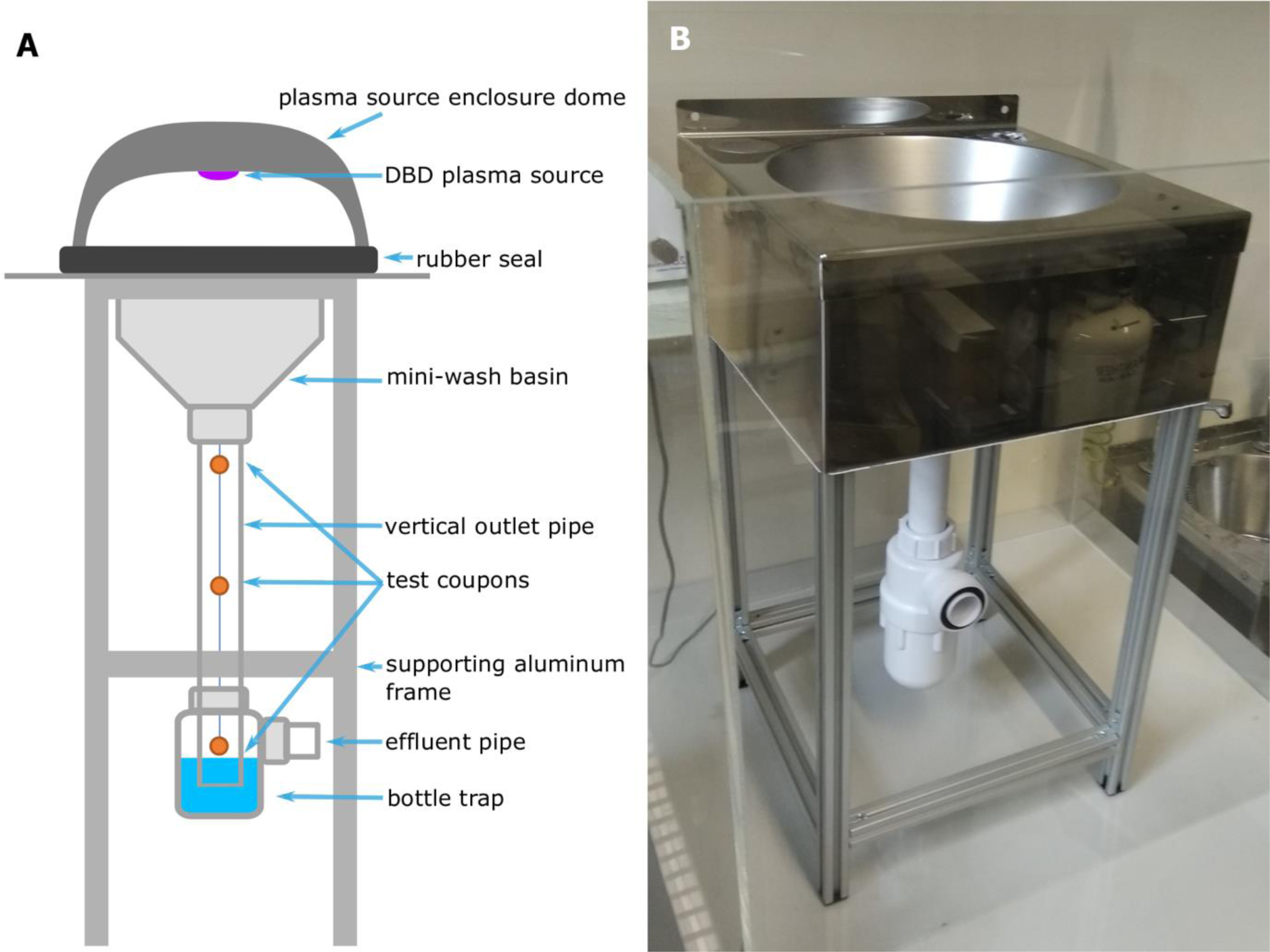
A) Schematic picture of the model wash basin with the DBD plasma source placed over the basin aperture and the placement of the coupons in the drain; B) a picture of a model wash basin enclosed in an acrylic tank separating the test rig from the rest of the laboratory.

## 4. Discussion

It is generally acknowledged that biofilms are more resistant to disinfection than planktonic cells; various mechanisms of this resistance have been proposed (Chapman, 2003; Donlan and Costerton, 2002; Hoff and Akin, 1986; Morita *et al*., 2015; Charron *et al*., 2023). Although studies exist showing the biocidal effect of gaseous ozone against planktonic and sporicidal bacteria, viruses and fungi (Broadwater *et al*., 1973; Dyas *et al*., 1983; Kowalski *et al*., 1998; Dev Kumar *et al*., 2020), its effects on biofilms are less known (Huth *et al*., 2009; Kaptan *et al*., 2014; Nicholas *et al*., 2013; Zargaran *et al*., 2017). The CDC bioreactor is an established tool used to generate repeatable biofilms in high-sheer conditions (Donlan and Costerton, 2002; Goeres *et al*., 2005; Gomes *et al*., 2014; Perez-Conesa *et al*., 2011), which are present, for example, in sink drains. This device is utilized in the ASTM E2562-12 standard (ASTM International, 2012), which describes a method of biofilm generation that furthermore uses *P. aeruginosa* – an opportunistic microorganism linked to many HAI outbreaks often connected to hospital water and waste systems, making it a relevant model to study.

The ability of ozone to inactivate microorganisms lies in its oxidising effect on the cell wall, particularly of lipids, and the induction of cell lysis (Thanomsub *et al*., 2002). Treating biofilms grown for up to 72h with gaseous ozone, we were unable to recover any culturable *P. aeruginosa*, and achieved a maximum log_10_ reduction of at least 7.4. A stricter control of the humidity in the experiment may yield higher log reduction values (Graves, 2012; Kogelschatz, 2003; Moisan *et al*., 2001; Zhang *et al*., 2016).

We explored the plasma-generated ozone disinfection efficacy by comparing it to SLD (Gasparini *et al*., 1995; Hernndez *et al*., 2000) and saw both to have equal efficacy when used on biofilms grown for 8 and 12h. Although this data suggests equivalent efficacy between ozone and SLD treatment, the application of ozone described offers several advantages, such as the ozone is manufactured, *in-situ* and on demand, from air. Furthermore, in complex real-life geometries where biofilms actually grow, such as water drainage systems, surface-tension restrictions mean that liquid disinfectants are unable to directly affect all areas of the target pipework. Generating gaseous ozone *in-situ* using the ambient air inside the pipework results in the gas permeating far more thoroughly than is possible with SLD.

Testing the efficacy of the plasma technology in more challenging conditions, by treating biofilm covered coupons placed in a model wash basin and drain assembly was also successful. Such experimental setups demonstrate the properties of plasma-generated ozone to penetrate small structures of the drain and successfully inactivate biofilm above the bottle-trap level, and most importantly, below the strainer or gasket; this reduces the risk of cross-contamination via back-splatter during use of the wash basin, which has been proven to pose an issue in healthcare facilities (Kotay *et al*., 2017, Garvey *et al.,* 2023). Preliminary results for ozone treatment of biofilm grown in a section of pipework show promising data that it is also possible to significantly reduce the bioburden of biofilms submerged in water (see Suppl. section 2.3). A recent work by Habimana and Casey showed that resident biofilm can promote the recruitment and incorporation of planktonic cells into an existing biofilm structure (Habimana and Casey, 2018). Hence, a further benefit of our approach is that plasma generated ozone treatment does not promote biofilm regrowth, in contrast to findings of Liang *et al*. (Liang *et al*., 2011).

The main advantage of this ozone generation technology lies in its ability to generate ROS from ambient air without the need for an external gas feed, and in such a way that the ozone region is safely isolated from the user. Since there is no requirement for feedstock gas, there is no risk of the sealed system under treatment being over-pressurised. This is particularly important in the drain context, since increasing the pressure could compromise the seal on the water trap. When treating a drain, the dome housing the plasma electrode creates a gas-tight seal around the drain inlet, with the water-trap itself forming the seal at the drain outlet. In this way, the treatment region is contained, and no ozone can leak to the outer environment (confirmed by ozone monitoring the immediate environment around the test drain). Furthermore, once the ozone treatment is concluded, and the ozone dissipates, there are no toxic residues left on the surface of the drain. As the ozone is generated electrically, from the ambient air, no hazardous chemicals need to be purchased or utilized; this is both cost effective and avoids contamination of wastewater, as well as subsequent oceanic pollution. Coupled with inherent consistency and ease of use, this technology makes a compelling alternative to existing techniques.

