## Supplementary Materials for "Antimicrobial efficacy of *in-situ* plasma-generated ozone against *Pseudomonas aeruginosa* biofilms in drains and water-submerged surfaces"

The content presented in this article is primarily concerned with validating the robustness of the method of biofilm recovery via sonication, as well as presenting preliminary data illustrating the efficacy of ozone disinfection of submerged biofilms grown on pipes; this also includes an evaluation of the robustness of the method for removing and quantifying biofilms from the corresponding pipework.

#### 1. Methods

##### 1.1 Growing biofilms on pipes

Three pieces of a PE-X pipe (British Standards Institution, 2007) (∅ 22 mm, 8.5 cm long) were connected using straight push-fit connectors with a snap-on end cap placed at the bottom end (Suppl. Fig. 4a). Standing upright with the end-cap at the bottom end, the assemble was filled to the second connector with the growth medium outlined in ISO-TS 15883-5:2005 Annex F (International Organization for Standarization, 2005). An overnight *P. aeruginosa* culture was diluted with fresh media and ca. 1 x $10^8$ CFU $ml^{-1}$ inoculated into the assembled pipework; this was incubated at 30°C for 3 days in a rotating incubator.

##### 1.2 Recovery of biofilms grown on pipes

After incubation, each 8.5 cm segment of tubing was disconnected and visually inspected for biofilm formation on the walls of the tube. The biofilm removal was adapted from ISO/TS 15883-5:2005 (International Organization for Standarization, 2005) as follows. Each segment was transferred to a 50 ml falcon tube with 10 ml ¼ strength Ringer's solution (Sigma-Aldrich, St. Louis, MO, USA) as a neutralizer, with 5g of sterilised natural sand. The tube was vortexed for 1 min and the sand allowed to settle. The resultant bacterial suspension was then diluted as required, plated on TSA plates, then incubated at 37°C overnight and enumerated. Mechanical disruption, as a method to remove biofilms that have formed on surfaces, has been shown to be an effective method, with sand grains being a sufficient size not to cause cell lysis (Gomes *et al.*, 2018).

#### 1.3 Experiments

##### 1.3.1 Validation of sonication recovery of biofilms

It is important gauge whether this method actually recovers bacteria in an effective manner, recapitulating expected growth trends. As such, this method was validated by quantifying biofilm growth on both polycarbonate and PTFE coupons at both room temperature (RT, 20-22°C) and 30°C. Biofilms were first grown on both types of coupon according to main text section 2.3, for 4 to 72h, and at the above temperatures. They were then removed from the bioreactor, and the bacterial growth quantified according to main text section 2.4. For each of the sampling times, temperature and type of coupon, at least 3 coupons were recovered. The limit of detection (LOD) was calculated based on the quantities and dilutions used. Results of this validation are presented in Suppl. Fig. 3.

##### 1.3.2 Validation of mechanical disruption recovery method

As above, it is important to evaluate the consistency of the recovery method. For biofilms grown as in section 1.1, the consistency of the biofilm formed on each segment was determined using the method described in section 1.2. This was done in triplicate.

##### 1.3.3 Ozone treatment of submerged biofilms

Two pipes with biofilm grown over the inner surface, treated as in section 1.1, were taken apart - after excess supernatant was discarded - and the segments bagged individually in generic polyethylene grip-seal bags, leaving some air inside before each bag was sealed. The diagram of the sample treatment is shown on Suppl. Fig. 4.b. One set of three segments was kept as an untreated control. One was kept dry and treated with one 15 s pulse of plasma. The remaining sections were submerged during treatment – either by replacing the end cap with a fresh one, placing the tube upright and filling it with 26.6 ml of ¼ strength Ringer's solution (vertical samples), or by filling the sampling bag with 126.6 ml of 5% Ringer's solution, ensuring the sample is fully submerged when treated (horizontal samples). The "wet" samples were treated with four 60 s pulses 10 min apart. All samples were left to dwell for 1h then the biofilm recovered as described in section 1.2.

To compare the efficacy of the ozone treatment, a control segment was exposed for 10 min to a control 1% solution of SLD as per manufacturer instruction.  The segment was then gently washed twice with sterile deionised water, and the bacteria were enumerated as described previously.

### 2. Results

#### 2.1 Validation of sonication recovery of biofilms

Suppl. Fig. 3 shows the number of recovered culturable *P. aeruginosa* from the polycarbonate and PTFE coupons. As expected, there were more colony forming units (CFU) recovered from the latter due to porousness of this material compared to the smoother polycarbonate. There were also more CFUs recovered from the biofilm grown at the 30°C than at the room temperature with the biggest difference observed at the earlier stages of the biofilm formation. Given the results followed the expected pattern and showed consistency, sonication was accepted as the recovery method and used in the following experiments. The LOD was estimated at 50 CFU $ml^{-1}$.

#### 2.2 Validation of mechanical disruption recovery method

The mean bacterial recovery from all nine pipe segments was $(3.7 \pm 0.4) \times 10^8$ CFU $ml^{-1}$. This highlights the consistency in the bacterial recoveries when biofilms were mechanically disrupted with sand. The LOD was estimated at 10 CFU $ml^{-1}$.

#### 2.3 Ozone treatment of submerged biofilms

The preliminary results show that the number of culturable *P. aeruginosa* recovered from a segment of a waste pipe treated with ozone, when dry, show at least 4.9 $log_{10}$ reduction compared to the untreated control, while the reduction for the sections submerged in water in horizontal and vertical position was at least 3.5 and 2.1 $log_{10}$, respectively. This data is presented in Suppl. Fig 5. Though no statistical significance can be quantified in this single-run experiment, the results are strongly indicative that ozone can be effective in treating biofilm in submerged pipes.

#### 2.4 Supplementary Figures

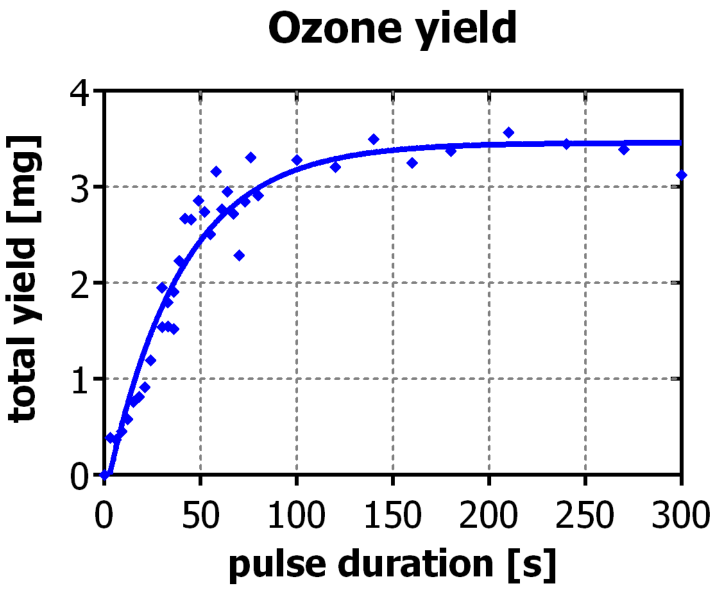


*Suppl. Fig. 1 Plasma source characteristics as a function of a pulse duration as measured using small, approx. 1-litre clear polyethylene grip-seal bags.*

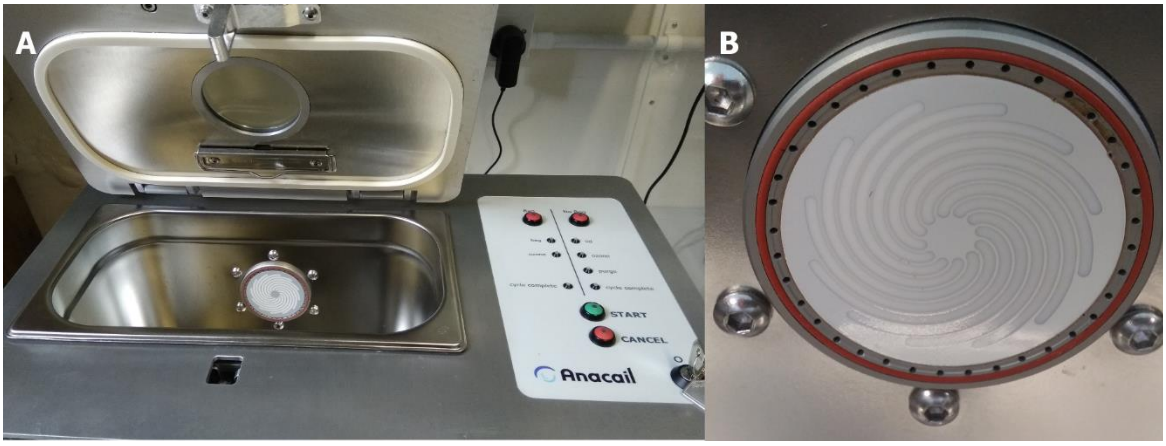


*Suppl. Fig. 2 A) prototype plasma-generation machine used to treat bagged samples; B) a close-up of the DBD plasma source*

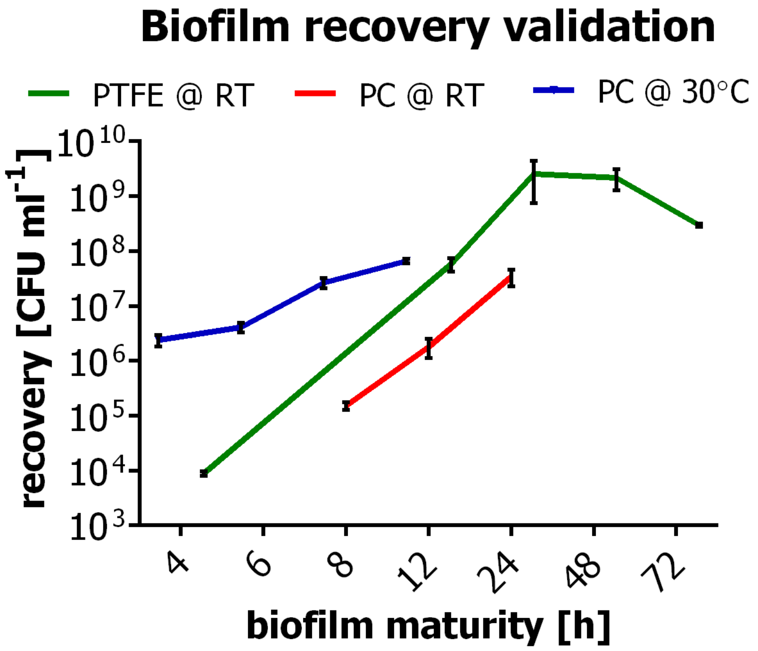


*Suppl. Fig. 3 Validation of the sonication as the recovery method of a biofilm grown for 4-24h on polycarbonate (PC) or PTFE coupons and at either room temperature (RT) or 30°C. Error bars show SEM.*

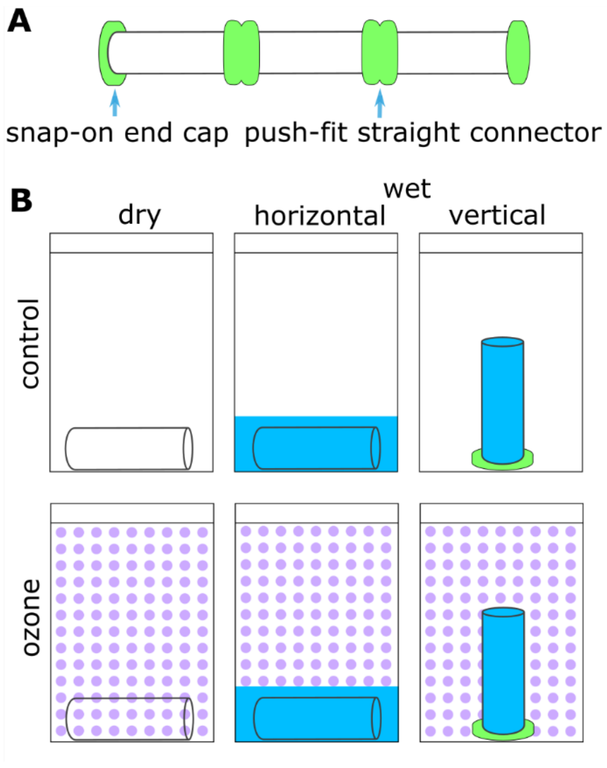


*Suppl. Fig. 4 A) Set up used for growing biofilm on the inside of the PE-X pipe; B) sample treatment*

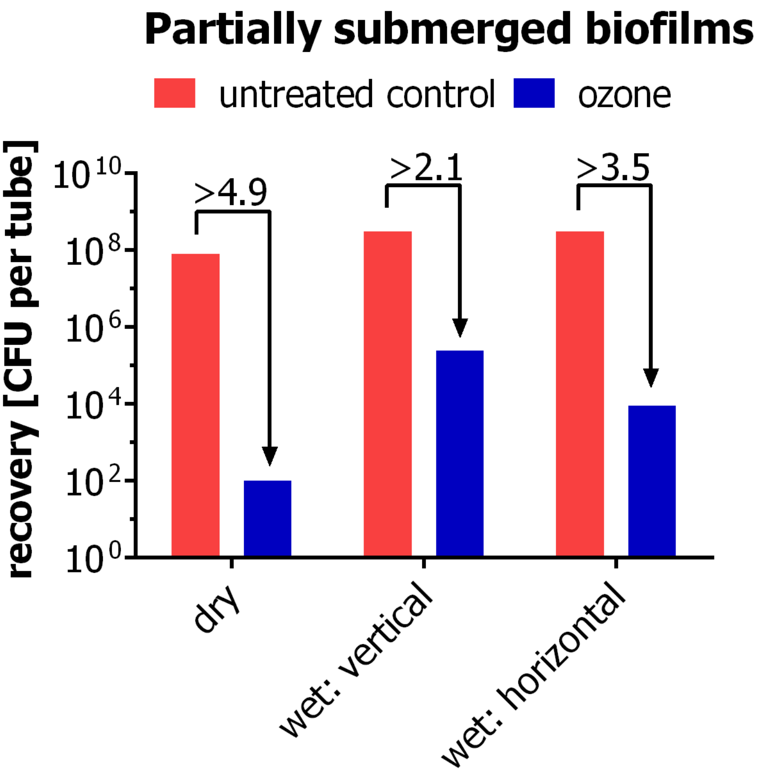


*Suppl. Fig. 5 Ozone treatment of dry and submerged biofilms grown on PE-X pipe*

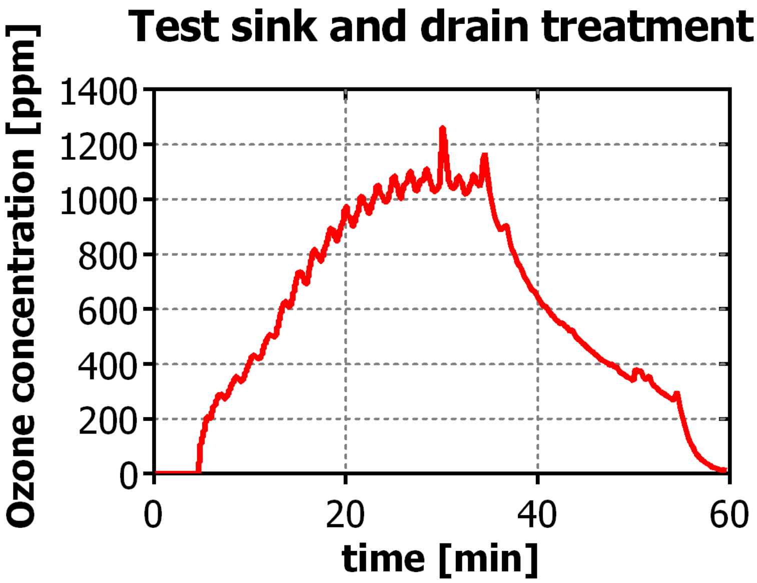


*Suppl. Fig. 6 30-min drain treatment protocol and dwell time used in the treatment of the model wash basin and drain assembly.*
